# Senolytic treatment targets aberrant p21-expression to restore liver regeneration in adult mice

**DOI:** 10.1101/759530

**Authors:** Birgit Ritschka, Tania Knauer-Meyer, Alba Mas, Jean-Luc Plassat, Daniel Sampaio Gonçalves, Hugues Jacobs, Elisa Pedone, Umberto Di Vicino, Maria Pia Cosma, William M. Keyes

**Author notes:** Correspondence to Bill Keyes, IGBMC, 1 Rue Laurent Fries, BP 10142, 67404 Illkirch - CU Strasbourg France.

## Abstract

Young mammals possess a limited regenerative capacity in tissues such as the liver, heart and limbs, but which is quickly lost upon maturation or transition to adulthood. Chronic cellular senescence is a known mediator of decreased tissue function in aging and disease. Here we investigated whether senescence plays a role in the progressive loss of liver regenerative capacity that develops in young adult mice. We find that following partial hepatectomy, the senescence markers p21, p16^Ink4a^ and p19^Arf^ become dynamically expressed at an age when regenerative capacity decreases. In addition, we demonstrate that treatment with a senescence-inhibiting drug improves regenerative capacity, through targeting of aberrant p21 expression. Surprisingly, we also find that the senescence marker p16^Ink4a^ is expressed in a different cell-population to p21, and is unaffected by senescence targeting. This work suggests that senescence may initially develop as a heterogeneous cellular response, and that treatment with senolytic drugs may aid in promoting organ regeneration.

## INTRODUCTION

Lower vertebrates retain the capacity to regenerate entire tissues, whereas mammals have a strikingly diminished regenerative capacity. Shortly after birth mammals are able to regenerate some organs, but this capacity declines rapidly, even before reaching adulthood (Bassat et al., 2017; Han et al., 2008; Payzin-Dogru and Whited, 2018; Porrello et al., 2011). The reasons for such loss of regeneration remain unclear.

The mammalian liver is one tissue that retains an increased regenerative capacity for a period of time, and therefore the model of two-thirds partial hepatectomy (PH) is frequently employed to study regeneration (Mitchell and Willenbring, 2008). Following surgery to remove 2/3 of the liver, the consequent tissue damage initiates a complex series of cellular events that results in the regrowth of the original liver mass over the course of about a week. Initially, damage associated cytokines and growth factors such as TNFa, and IL6 are released rapidly after the resection (Michalopoulos, 2007; Su et al., 2002; Yamada et al., 1997). This sets in motion a cascade where the different populations of liver cells coordinate and contribute to the regeneration of the tissue (Michalopoulos, 2007; Pedone et al., 2017). However, while this process is very effective in young animals, there is a rapid decline in the efficiency of regeneration, and adult mice have a significantly reduced capacity to regenerate, and present with increasing liver damage and mortality following PH (Enkhbold et al., 2015; Loforese et al., 2017; Mitchell and Willenbring, 2008).

Cellular senescence is a form of permanent cell-cycle arrest linked with aging and tissue damage (Childs et al., 2015; Munoz-Espin and Serrano, 2014). The aberrant accumulation of senescent cells expressing markers such as p16^Ink4a^, p19^Arf^ and p21, and secreting inflammatory factors of the “senescence-associated secretory phenotype” (SASP), contributes to the loss of tissue function seen during chronological aging and in many diseased states (Baker et al., 2016; Baker et al., 2011; Childs et al., 2017; Munoz-Espin and Serrano, 2014). This perhaps is best exemplified with mouse models in which p16-positive cells are genetically eliminated, thus resulting in increased lifespan and healthspan (Baker et al., 2016; Baker et al., 2011), demonstrating that chronic senescence is detrimental for tissue function. More recently, the elimination of senescent cells through pharmacological means has also been demonstrated to delay the aging process and ameliorate disease symptoms. Interestingly, several drugs that target senescent cells for elimination, collectively known as “senolytics”, have been successfully used to increase lifespan, healthspan, and to improve some disease-associated conditions, including Alzheimer’s, Parkinson’s, atherosclerosis and liver steatosis (Baar et al., 2017; Bussian et al., 2018; Chang et al., 2016; Chinta et al., 2018; Musi et al., 2018; Ogrodnik et al., 2017; Xu et al., 2018; Yosef et al., 2016). While these drugs can target the apoptotic machinery to kill senescent cells, in many cases, their precise cellular or molecular targets in vivo remain unclear.

Here, using the model of 2/3 PH, we investigated whether the decrease in regenerative capacity that develops in adult animals is linked to senescence, and whether this could be improved with senolytic drug treatment.

## RESULTS

To investigate a possible connection between senescence induction and early loss of regenerative capacity, we performed 2/3 PH in young (2-3 months) and adult (6-8 months) mice and analyzed regeneration in each group during one week after surgery. The livers in young mice regenerated, reaching almost original liver/body weight ratio by seven days after PH (Figure 1A). However, the livers of the adult mice presented with a significant decrease in regeneration (Figure 1A). Assessment of the histology in young mice (Figure 1B) identified regeneration as previously described, without necrosis or inflammation (Mitchell and Willenbring, 2008). The adult livers after PH showed signs of steatosis, particularly prolonged micro- and macro-vacuolization (Figure 1B, day 3). This is similar to previous reports that have shown that aged livers (>12 months with PH, or >18 months without PH) displayed marked steatosis (Loforese et al., 2017; Ogrodnik et al., 2017). This suggests that injury such as PH in adult mice, induces changes that are normally seen during aging.

**Figure 1:**
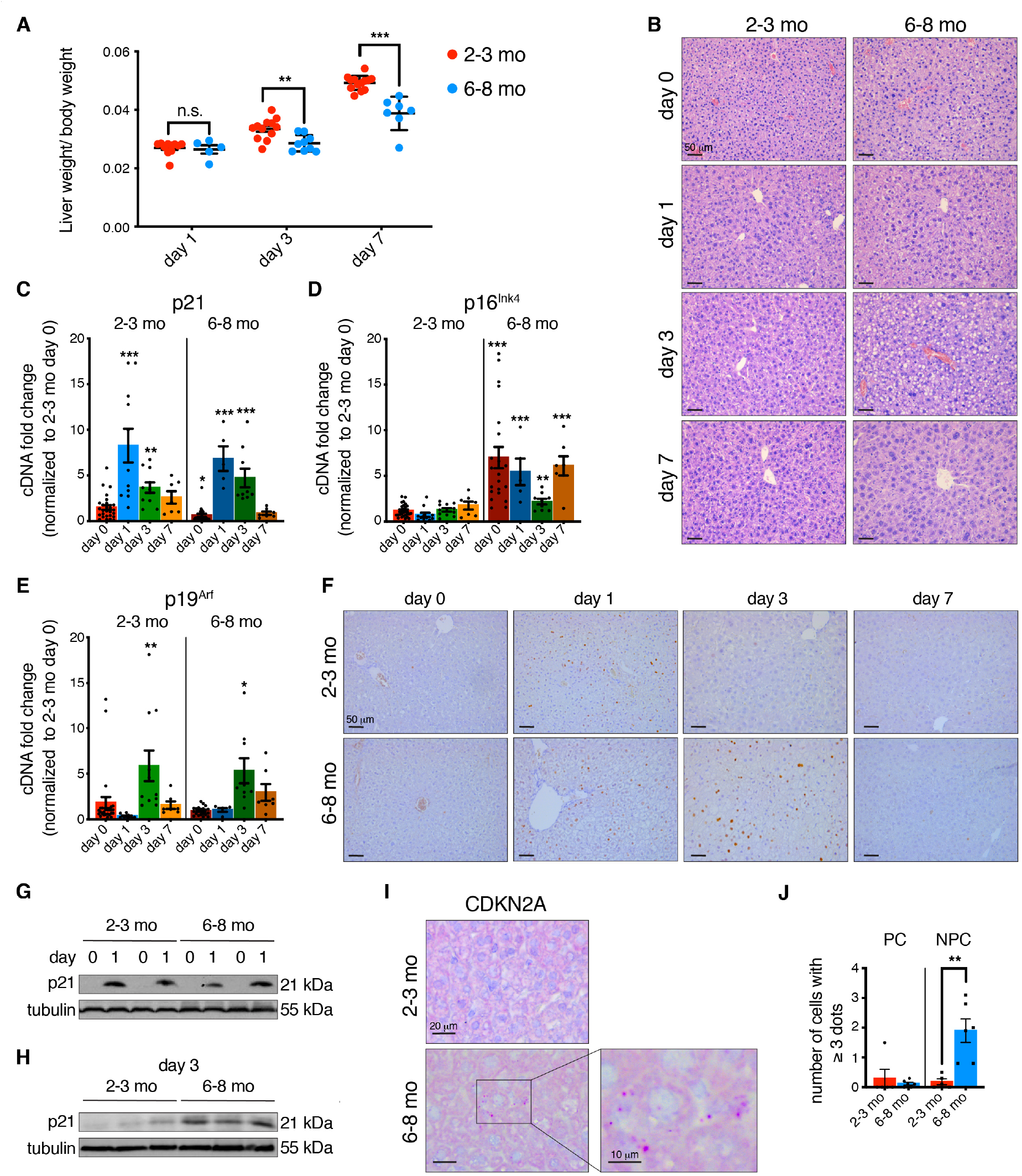
p21 exhibits prolonged expression in adult livers with impaired regeneration. (A) Liver regeneration of young (2-3 months) and adult (6-8 months) mice was calculated as liver/body weight ratio at different days after PH (n=5-12). (B) H&E staining of young and adult liver sections before (day 0) and at different time points after PH. All images are representative of at least 5 biological replicates. Scale bars 50 μm. (C-E) qPCR analysis for the senescence markers p21 (C), p16^Ink4a^ (D) and p19^Arf^ (E) of young and adult livers at different days after PH normalized to young livers at day 0 (n=5-27). (F) Immunohistochemistry for p21 in young and adult liver sections at different days after PH. All images are representative of at least 3 biological replicates. Scale bars 50 μm. (G-H) p21 expression in whole liver lysates at (G) 0 and 1 and (H) 3 days after PH measured by western blot. Tubulin was used as a loading control (day 0 and 1: n=2, day 3: n=3). (I) RNA in situ hybridization staining for CDKN2A in young and adult liver sections before PH (day 0). Scale bars 20 μm. Boxed area shows higher magnification of positive staining. Scale bar 10 μm. All images are representative of at least 5 biological replicates. (J) Quantification of RNA in situ hybridization staining for CDKN2A shows number of cells/field of view that have ≥ 3 dots/cell. PC = parenchymal cells, NPC = non-parenchymal cells (2-3 mo: n=5, 6-8 mo: n=6). Error bars, mean ± SEM, unpaired two-tailed Student’s *t*-test (*p ≤ 0.05, **p ≤ 0.01, ***p ≤ 0.001).

Given that adult mice exhibit decreased regeneration by 6-8 months, we assessed for expression of core senescence markers p21, p16^Ink4a^ and p19^Arf^ by qPCR in the young and adult livers. In pre-hepatectomy liver samples (day 0), we found that baseline levels of *p21* decreased between 2-3 months and 6-8 months of age (Figure 1C). Conversely however, the senescence marker *p16^Ink4a^* significantly increased during this timeframe, while there was no change in *p19^Arf^* expression (Figure 1D-E). This suggests that a prominent age-associated senescence marker is induced in adult mice during this time window. Previously, the induction of senescence and p16^Ink4a^ in the liver was linked with fibrosis (Krizhanovsky et al., 2008). Therefore, we checked for alterations in the levels of fibrosis, which is negligible in young mice following PH. However, no significant change was seen in adult animals, either before or following PH (Figure S1A).

Next, we assessed for dynamic changes in the expression of the same core senescence markers during the phase following PH, both in young and adult mice. As previously described, following PH in young mice, there was a transient, p53-independent increase in *p21* expression which peaked about 24-48 hours post-PH and which decreased by day 7 (Figure 1C) (Albrecht et al., 1997; Buitrago-Molina et al., 2013; Stepniak et al., 2006). In older animals, this induction of *p21* also occurred (Figure 1C). Surprisingly, similar analysis for *p16^Ink4a^* revealed that even though expression was increased in adult pre-hepatectomy livers, this did not increase further, and even decreased after PH, before returning to the elevated adult pre-hepatectomy level (Figure 1D). Finally, *p19^Arf^* expression peaked transiently at day 3 post-PH in both young and adult animals (Figure 1E).

Based on these findings, we next examined the expression pattern and localization of p21 at the protein level. In agreement with our qPCR results and previous reports, there was a clear transient induction of p21 one day after PH in young animals, predominantly in hepatocytes (Figure 1F-G). Surprisingly however, while a similar induction was seen 24 hours after PH in adult livers, p21 expression persisted longer, now also being detectable at day 3 after PH, when p21 expression is lost in the young (Figure 1F). Comparison of the total protein level in young and adult tissues three days after PH by western blotting showed clearly that p21 levels remain aberrantly high in mice at 6-8 months of age (Figure 1H).

Our qPCR data also showed a significant increase in the senescence mediator p16^Ink4a^ prior to PH. To determine where p16^Ink4a^ is expressed, we performed in situ hybridization using a probe that recognizes a shared exon in the *CDKN2A* locus, common to both *p16^Ink4a^* and *p19^Arf^*. As our qPCR results showed that there is no expression of *p19^Arf^* in adult tissue before hepatectomy, the observed signal in the adult livers corresponds to *p16^Ink4a^* expression (Figure 1I-J and S1B). Similar results were obtained with a lower affinity probe specific for *p16^Ink4a^* (data not shown). Surprisingly however, the staining with the CDKN2A probe clearly showed that *p16^Ink4a^* is not in the hepatocytes, and is more likely in the non-parenchymal stellate, Kupffer or endothelial cells (Figure 1I-J) (Demaria et al., 2014; Krizhanovsky et al., 2008; Liu et al., 2019). In addition, we used this same probe to examine the *p19^Arf^* expression that was detected in young mice at day 3, when *p16^Ink4a^* was not detectable, and find that this was also not localized in the hepatocytes (Figure S1C).

Together, these data show that there are dynamic changes in core senescence mediators in adult liver that has an impaired capacity to regenerate. Senescent cells are also associated with an increased inflammatory environment in vivo, and exhibit a pronounced secretory capacity, known as the SASP. To determine whether there is a mis-expression of SASP factors before or after PH in adult tissues, we next examined for their expression. We checked by qPCR whether common SASP factors such as *IL6, Ccl2, IL1* or *Pai1* are similarly increased prior to PH in adult tissue. However, none of the SASP factors examined were increased in the pre-hepatectomy adult livers (Figure S1D). Our data pointed to an aberrant expression of p21 three days after PH in adult compared to young tissue. Based on this, we analyzed the expression of 111 proteins, including many cytokines, chemokines and inflammatory factors that are common to the SASP. Interestingly, we found that many of these showed increased expression in day 3 adult livers compared to their young counterparts (Figures 2A-C and S2A). These included the SASP and liver damage factor IGFBP1, as well as Ccl2, Csf1, Pai-1 and Cxcl2. This demonstrates that the adult tissue has increased expression of many SASP factors following PH in adult mice.

**Figure 2:**
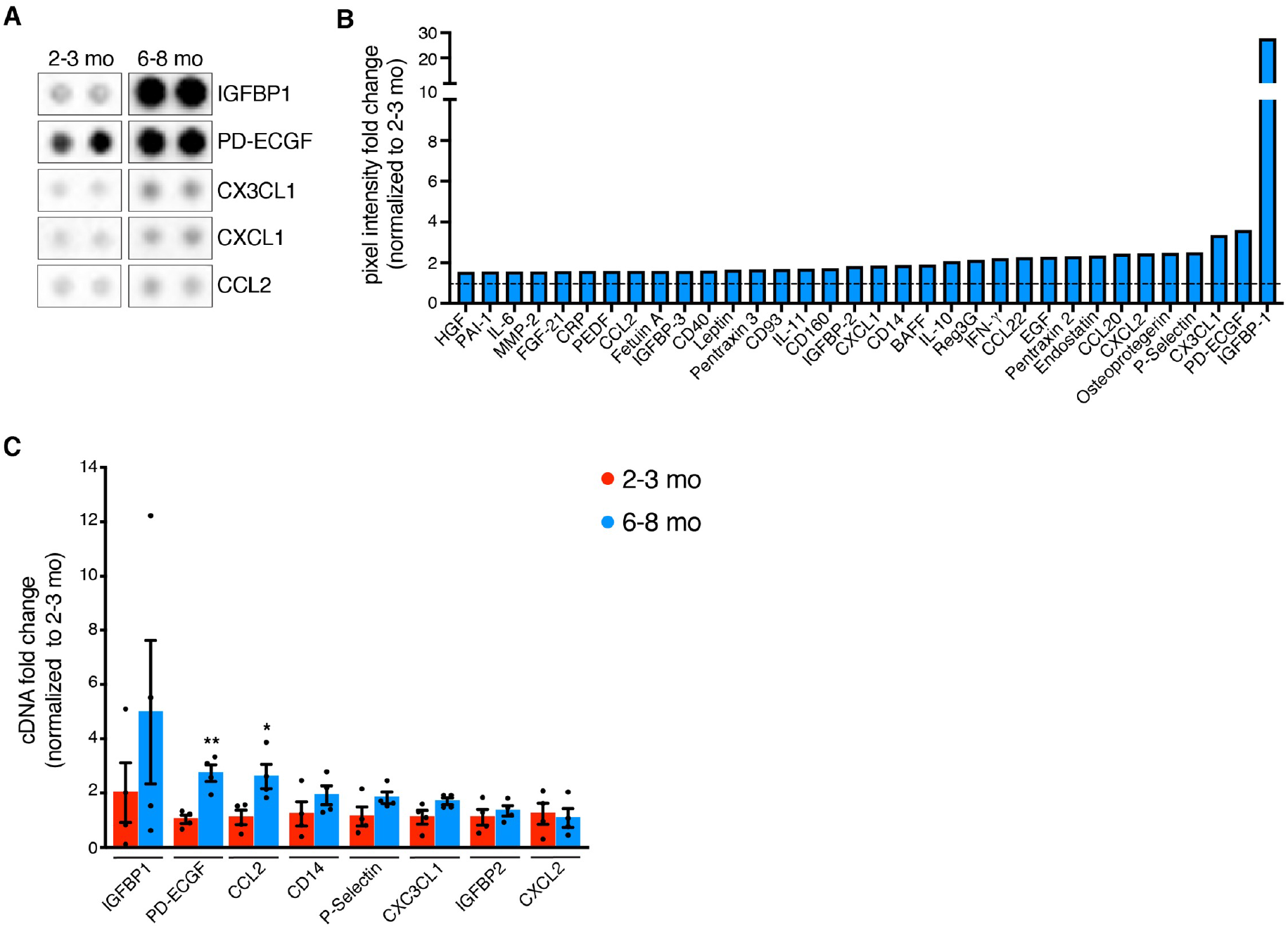
Adult non-regenerating livers show an increase in many SASP proteins. (A) Representative images from cytokine array membranes of selected cytokines of 2-3 and 6-8 months old WT livers 3 days after PH. All cytokines are detected in duplicates. (B) Relative expression ratio (≥1.5 fold) of 6-8 months old liver tissue 3 days after PH normalized to 2-3 months old livers (n=2). (C) qPCR analysis for some upregulated cytokines in 6-8 months old livers 3 days after PH normalized to 2-3 months old livers (n=4). Error bars, mean ± SEM, unpaired two-tailed Student’s *t*-test (*p ≤ 0.05, **p ≤ 0.01, ***p ≤ 0.001).

Together, our data suggests that there is an associated increase in senescence mediators when the adult liver loses its capacity to regenerate. We next asked if treatment with a senolytic drug could restore or improve regenerative capacity in adult livers. To test this hypothesis, we devised a strategy to pretreat livers with the senolytic compound ABT-737, or vehicle control, twice over two days immediately prior to PH, as such delivery could potentially target both p16^Ink4a^- and p21-positive cells (Figure 3A). When we measured regeneration, adult mice of 6-8 months treated with vehicle alone showed decreased regenerative function, similar to their untreated controls (Figure 3B-C). Surprisingly however, mice that were pre-treated with ABT-737 showed significantly increased regenerative capacity by day 7 after hepatectomy (Figures 3B-C). To support this, we assessed if liver function was also improved, and measured serum levels of the liver enzymes aspartate transaminase (AST) and alanine transaminase (ALT), which are increased upon PH, tissue damage and aging (Enkhbold et al., 2015; Loforese et al., 2017; Raven et al., 2017). This further supported that senolytic-treated livers were functionally improved, as they exhibited significantly reduced AST levels (Figure 3D). Improvements were also evident at the histological level. Treatment with DMSO-vehicle alone further exaggerated the phenotype of adult mice, resulting in increased steatosis and more mitotic figures (Figure 3E). However, the ABT-737 treated tissue appeared more like the young counterpart (Figure 3E). Together, this demonstrates that pre-treatment of liver with a senolytic compound improves regenerative capacity.

**Figure 3:**
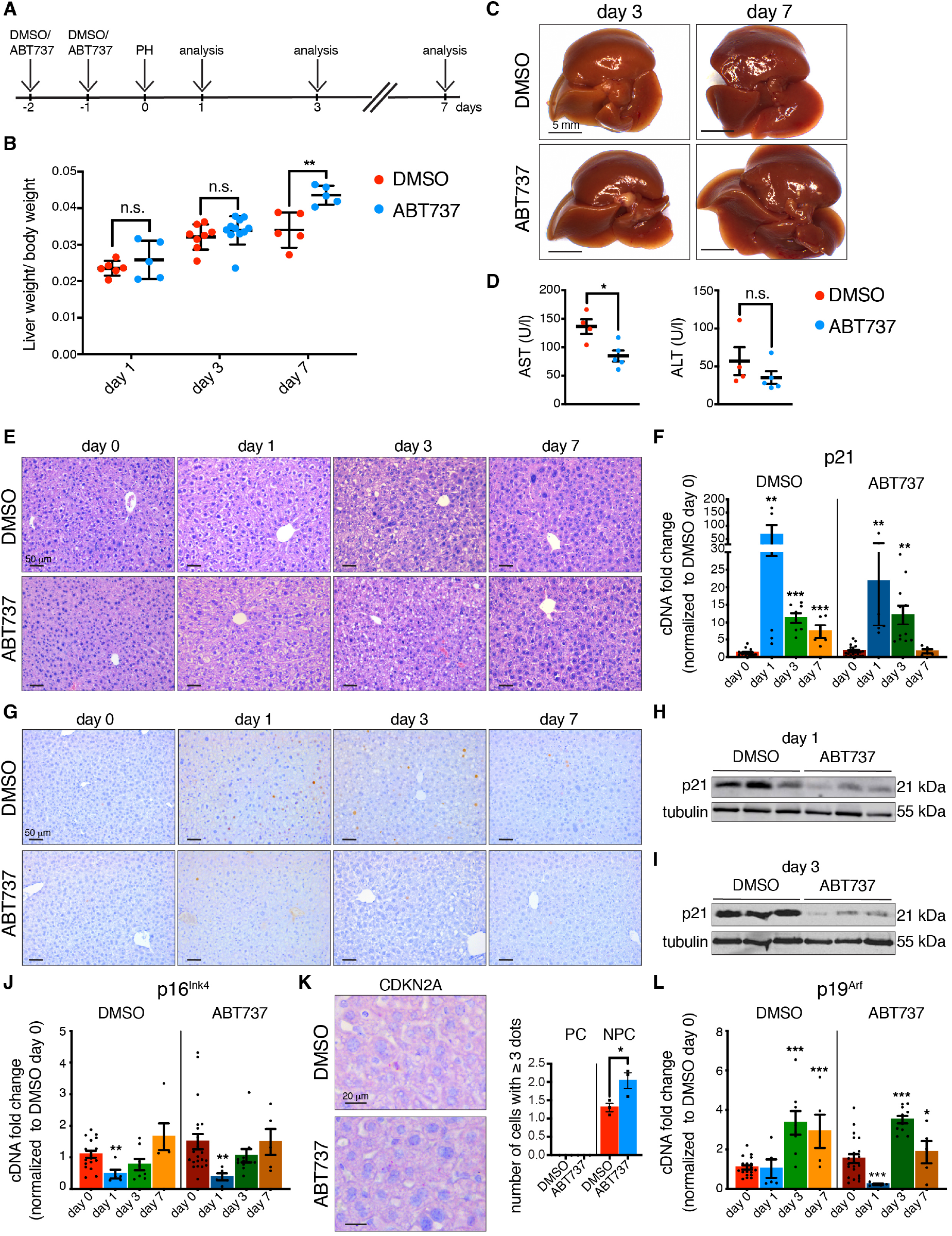
ABT-737 improves regenerative capacity in adult livers. (A) Schematic representation of the experimental protocol. Intra-peritoneal injection of DMSO or ABT-737 twice over 2 days was followed 1 day later by PH and mice were analyzed 1, 3 and 7 days after PH. (B) Liver/body weight ratios of 6-8 months DMSO or ABT-737 treated mice at different time points after PH (n=5-11). (C) Representative macroscopic images of livers from DMSO or ABT-737 treated mice 3 and 7 days after PH. Scale bars 5 mm. (D) Aspartate aminotransferase (AST) and alanine aminotransferase (ALT) measured 7 days after PH (DMSO: n=4, ABT-737: n=5). (E) H&E staining of DMSO and ABT-737 treated livers before and at different days after PH. All images are representative of at least 5 biological replicates. Scale bars 50 μm. (F) qPCR analysis for p21 in DMSO and ABT-737 treated livers at different days after PH, normalized to DMSO day 0 (n=5-21). (G) Immunohistochemistry for p21 in DMSO and ABT-737 treated livers at different time points after PH. All images are representative of at least 4 biological replicates. Scale bars 50 μm. (H-I) p21 expression following treatment with DMSO or ABT-737, in whole liver lysates 1 (H) and 3 (I) days after PH, measured by western blot. Tubulin was used as a loading control (n=3). (J) qPCR analysis for p16^Ink4a^ in DMSO and ABT-737 treated livers at different days after PH normalized to DMSO day 0 (n=5-21). (K) RNA in situ hybridization staining and quantification for CDKN2A in DMSO and ABT-737 liver sections before PH (day 0). All images are representative of at least 3 biological replicates. Scale bars 20 μm. Quantification shows number of cells/field of view that have ≥ 3 dots/cell. (n=3) PC = parenchymal cells, NPC = non-parenchymal cells. (L) qPCR analysis for p19^Arf^ in DMSO and ABT-737 treated livers at different days after PH normalized to DMSO day 0 (n=5-21). Error bars, mean ± SEM, unpaired two-tailed Student’s *t*-test (*p ≤ 0.05, **p ≤ 0.01, ***p ≤ 0.001).

To explore the mechanisms by which this senolytic exerts its beneficial effects, we examined the expression patterns of the core senescence mediators following treatment. First, examination of the *p21* transcript by qPCR revealed *p21* levels were decreased at day 1 post PH in the senolytic versus vehicle samples (Figure 3F). Furthermore, when we examined for p21 by immunostaining, it was evident that there was a significant reduction in p21 expression, with most senolytic-treated hepatocytes now staining negative for p21 (Figure 3G). This was most evident when we performed western blot on whole liver lysates, which demonstrated an overall decrease in p21 expression one and three days after PH (Figure 3H-I). Next, we examined for *p16^Ink4a^* expression. Interestingly, while p16^Ink4a^ is described as a primary mediator of aging-associated tissue decline, and p16^Ink4a^-positive cells are suggested as a primary target of senolytic treatment in aged and damaged tissues, here we found no change in *p16^Ink4a^* transcript levels upon senolytic treatment (Figure 3J). In addition, the distribution of *p16^Ink4a^* as assessed by in situ hybridization was unchanged (Figures 3K and S1E), suggesting that these cells are not affected by senolytic treatment, nor that they significantly contribute to the loss of regenerative capacity in these tissues at this age. Next, we examined for changes in *p19^Arf^* expression. While treatment with senolytic appears to decrease the expression of *p19^Arf^* on day 1 after PH, the levels at days 3 and 7 remain unchanged (Figure 3L). Altogether, this data suggests that the regenerative capacity of the liver can be restored by treatment with senescence-targeting drugs.

Our results imply that mis-expression of p21 is the major contributor to the loss of liver regeneration that develops in adult mice. To confirm this, we performed PH in mice deficient for p21 at both young and adult ages. At 2-3 months, p21-deficient mice displayed similar regeneration as their young wildtype controls at day 3, but presented with a delay in regeneration by day 7 (Figure 4A). However, when compared to age-matched wildtype controls, adult p21-deficient mice now showed significantly increased regenerative capacity at day 3 post-PH, and with continued growth to day 7 (Figure 4A). Histological analysis revealed no major differences in comparison to age-matched controls (Fig 4B). Furthermore, by qPCR analysis, we found that p21-deficient mice also had similar expression patterns of *p16^Ink4a^* and *p19^Arf^* following PH, as was seen in wildtype mice (Figure 4C-D).

**Figure 4:**
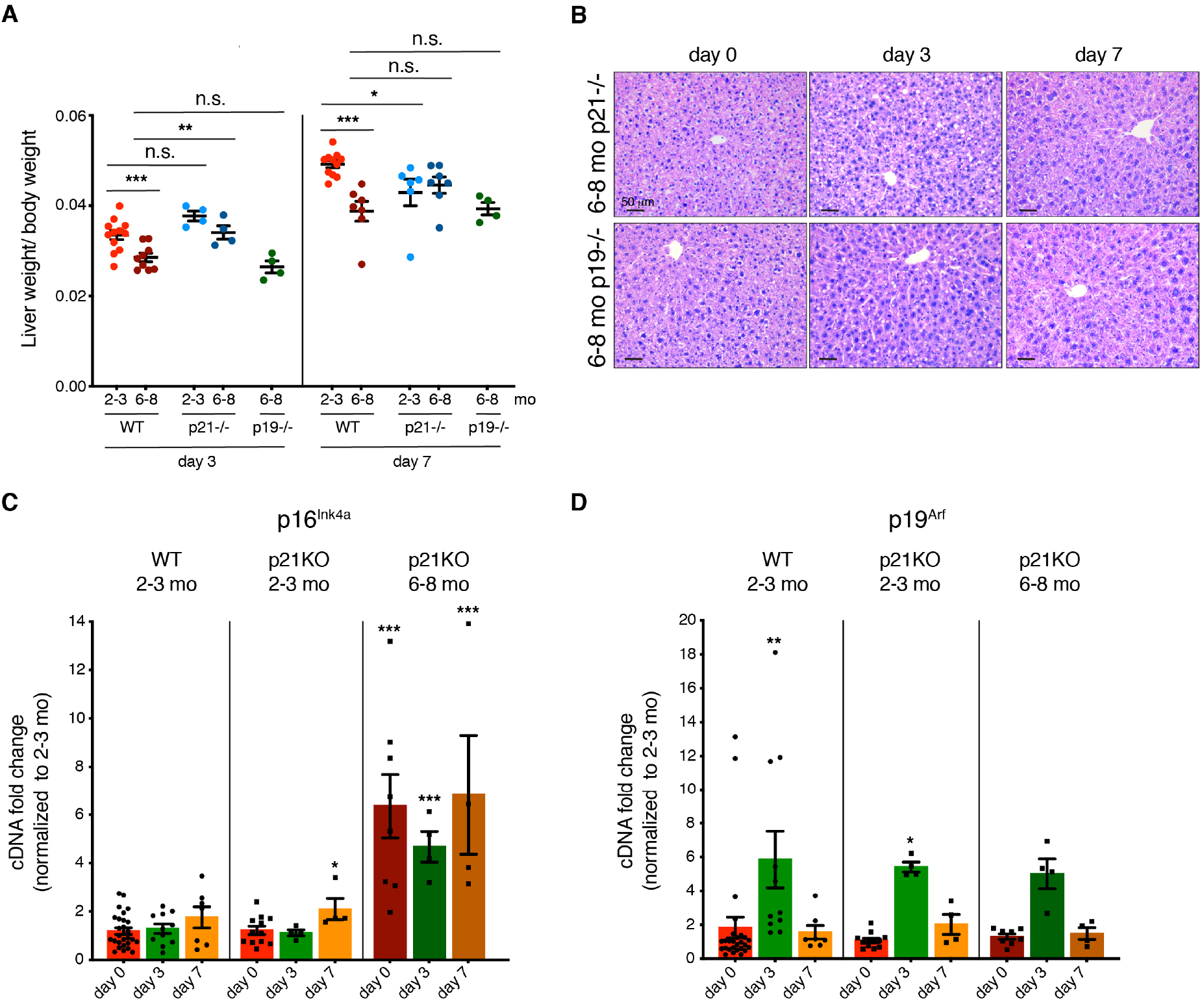
Adult p21-deficient mice have improved regenerative capacity. (A) Liver/body weight ratios of 2-3 and 6-8 months *WT, p21*-/- and *p19*-/- mice at 3 and 7 days after PH (n=4-12). (B) H&E staining of 6-8 months old *p21*-/- and *p19*-/- livers before and at different days after PH. All images are representative of at least 4 biological replicates. Scale bars 50 μm. (C-D) qPCR analysis for (C) p16^Ink4a^ and (D) p19^Arf^ of 2-3 months old WT and p21-/- and 6-8 months old p21-/- livers at different days after PH normalized to 2-3 months old WT day 0. (n=4-27).

Our data also uncovered a transient increase in p19^Arf^ expression following PH. To investigate whether this is functionally involved, we performed PH on p19^Arf^-deficient mice at 6-8 months of age. However, loss of p19^Arf^ had no effect on regeneration capacity and these mice performed as the adult wildtype controls (Figure 4A-B). Together, these data demonstrate that the aberrant prolonged expression of p21 becomes a contributory factor in the decreased regeneration of this tissue in adult mice.

## DISCUSSION

Here we uncover that p21 becomes aberrantly expressed following PH in adult mice, which contributes to the age-associated loss of regenerative capacity in adult liver. Surprisingly, this appears independent of other senescence mediators p16^Ink4a^ and p19^Arf^. We show that interference with this process by pharmacological means can improve functional regenerative capacity. This suggests that such an approach could be developed to aid in problems associated with decreased regeneration, for example to improve organ transplant efficiency or regeneration after liver surgery.

Many tissues, including the liver, heart and limbs possess a limited regenerative capacity in newborn or young mice, but which is lost upon maturation. As far as we are aware, mis-regulation of senescence has not been causally linked to such loss of regenerative capacity. Here, we first assessed the dynamic patterns of senescence markers during liver regeneration in young and adult mice. Although a transient p53-independent increase in p21 is well described following PH, its precise functions remain unclear, and loss of p21 does not seem to adversely impact regeneration in young animals. However, in models of advanced aging and severe liver damage, aberrantly expressed senescence markers, including p21 and p16^Ink4a^ have been reported to impede liver regeneration. For example, in models of liver fibrosis induced by CCl4 treatment, a robust senescence response is induced primarily in the stellate cells, which serves to limit fibrosis (Krizhanovsky et al., 2008). Other models of severe liver damage such as extended hepatectomy (Lehmann et al., 2012), acetaminophen treatment (Bird et al., 2018), Mdm2-deletion (Lu et al., 2015), β1-integrin loss ((Raven et al., 2017) or p21-overexpression (Raven et al., 2017), each induce a pronounced p21-expression in hepatocytes, which results in decreased regeneration, senescence, and senescence-spreading. It will be interesting to determine if senolytic-treatment can also improve such cases of severe damage. Indeed, senolytics have already been shown to function in the liver of mouse models with immune dysfunction (Ovadya et al., 2018), and in ameliorating age-related hepatic steatosis (Ogrodnik et al., 2017). Interestingly, as PH is associated with induction of steatosis in older animals as shown here and elsewhere (Loforese et al., 2017), our study further supports that these drugs could aid in liver regeneration in elderly patients.

Senolytic treatment is increasingly shown to have beneficial effects in enhancing tissue function and alleviating disease symptoms in a variety of tissues (Childs et al., 2017; Xu et al., 2018). However, in many cases, the specific cellular targets or molecular mediators in vivo remain to be identified. Our study suggests that p21-positive cells may be a primary target. This is supported by the findings that protection from apoptosis is a main function of p21, including in senescent cells (Storer et al., 2013; Yosef et al., 2017), and many senolytics, including the one used here, work by blocking anti-apoptotic pathways (Yosef et al., 2016). In addition, as p21 functions to protect cells from damage, prolonged loss of p21 in aging mice predisposes to cancer through loss of this cytoprotective effect (Martin-Caballero et al., 2001). Interestingly, our study suggests that a one-time removal of p21 positive cells using a senolytic has a beneficial effect on regeneration, but probably without the long-term consequences of p21-loss.

Surprisingly however, we see no effect of senolytic treatment on the increased expression of p16^Ink4a^ that is present prior to hepatectomy, and which becomes detectable at the same stage as the decrease in regenerative capacity. Many studies show how targeting p16^Ink4a^ expression has beneficial effects on aged and damaged tissue (Baker et al., 2016; Baker et al., 2011). However, in most cases, this also results in reduction of p21 and p19^Arf^, making it difficult to discern specific effects of each gene. Furthermore, in some contexts, it appears p16^Ink4a^ is required for beneficial effects including during wound repair, pancreatic function during aging and in vivo reprogramming (Demaria et al., 2014; Helman et al., 2016; Mosteiro et al., 2016), the latter being another setting where p21 expression is detrimental (Mosteiro et al., 2018). As p16^Ink4a^ and p21 are expressed in different cell populations in our study, this hints that p16^Ink4a^, at this level of expression at least, may have beneficial effects in the liver also. However, why senolytic treatment seems to eliminate p16^Ink4a^ positive cells in other contexts and not here remains unknown, but probably relates to the level of p16^Ink4a^ - expression or co-expression with other senescence genes, as p16^Ink4a^ levels become increasingly higher with age (Burd et al., 2013; Krishnamurthy et al., 2004). Perhaps with advanced age or chronic damage, p21 and p16^Ink4a^ become co-expressed, and at higher levels in the same cell types (Wang et al., 2014), resulting in a full-senescence response, and what we witness here is an early stage in a cumulative and progressive decline that becomes more complex over time.

## MATERIALS AND METHODS

### Animal use

Wild-type C57Bl6/J, *p21*-/- and *p19*-/- were maintained in a temperature- and humidity-controlled animal facility with a 12h light/dark cycle. Breeding and maintenance of mice were performed in the accredited IGBMC/ICS animal house, in compliance with French and EU regulations on the use of laboratory animals for research, under the supervision of Dr. Bill Keyes who holds animal experimentation authorizations from the French Ministry of Agriculture and Fisheries. All animal experiments were approved by the Ethical committee Com’Eth (Comité d’Ethique pour l’Expérimentation Animale, Strasbourg, France).

### Partial hepatectomy (PH)

The mice used in this study were 8-14 weeks (young) and 24-32 weeks (adult) male and female mice. Two-thirds partial hepatectomy was performed under isoflurane anesthesia and according to a standard procedure (Mitchell and Willenbring, 2008). In brief, after opening of the abdomen, the median and left lateral lobes were ligated and removed. Liver samples were collected during PH (day 0-time point) and at indicated time points following the surgery. For measuring the weight of the regenerating liver following surgery, only the nonresected lobes were measured. For RNA and protein analysis, liver samples were snap-frozen in liquid nitrogen. For histological and RNA *in situ* analysis, livers were fixed with 10% neutral buffered formalin for 24 hrs at 4°C and at room temperature, respectively.

### Elimination of senescent cells *in vivo*

For elimination of senescent cells, 6-8 month old mice were injected once per day intra-peritoneally (i.p.) on 2 consecutive days immediately before PH with DMSO (2%) or ABT-737 (25 mg/kg body weight; AdooQ Bioscience). DMSO and ABT-737 were prepared in the following working solution: 30% propylene glycol, 5% Tween 80, 5% dextrose in water (pH 4-5) as previously reported (Ovadya et al., 2018).

### Serum collection and analysis

Mice were anesthetized by isoflurane inhalation and blood was taken by retro-orbital bleeding. Mice were immediately euthanized afterwards by CO_2_ inhalation. The tests are performed with an AU-400 automated laboratory work station (Beckman Coulter France SAS, Villepint, France) at the Institut Clinique de la Souris.

### Histology and immunohistochemistry

Fixed liver tissues were washed in PBS and then processed for paraffin embedding and hematoxylin and eosin (H&E) staining. Immunohistochemistry was performed using standard procedures. Antigen retrieval was performed by boiling deparaffinised sections for 30 min in Tris-EDTA at pH 9.0 in a pressure cooker. Primary antibodies were incubated overnight at 4°C, and secondary antibodies were incubated for 1 hr at room temperature in 1% serum, 0.1% Triton-X in PBS. Sections were treated with streptavidin-biotin-peroxidase (Vectastain Elite, Vector, Burlingame, Calif.) and immune complexes were visualized with diaminobenzidine (Vector, Burlingame, Calif.) as per manufacturer’s protocol. Primary antibodies were used at the following dilutions: anti-p21 (1:50, HUGO-291, CNIO). Biotin-conjugated secondary antibodies (Vectastain Elite) were used at a dilution of 1:2000. Representative pictures were acquired with a Leica DM 1000 LED microscope.

### Sirius red staining

Sirus red staining was performed according to standard protocols. Histological sections were digitalized using a digital slide scanner (Nanozoomer HT 2.0; *Hamamatsu*) and then scored for fibrosis using imageJ software and dedicated homemade macro. Results were analyzed using 2-Way ANOVA (Prism 6.04; *GraphPad*).

### RNA *in situ* hybridization

In situ RNA hybridization was performed using RNAscope probes for CDKN2A, PPIB positive control and bacterial Dapb negative control (catalog no’s. 411011, 313911 and 310043, Advanced Cell Diagnostics) as per the manufacturer’s instructions. RNAscope 2.5 HD Reagent kit-RED was used for chromogenic labelling and hematoxylin for counterstaining. Images were obtained on a Leica DM 1000 LED microscope.

### Western blot

Frozen tissue was homogenized in the following lysis buffer: 50 mM Tris/HCl (pH 7.5), 1 mM EDTA, 1 mM EGTA, 1% v/v Triton X-100, 1 mM Na3VO4, 50 mM NaF, 5 mM Na_4_P_2_O_7_×10H_2_O, 0.27 M Sucrose in MiliQ H_2_O with freshly added 1 mM DTT, 0.2 mM PMSF and 1X Protease Inhibitor Cocktail (Roche). Lysates were cleared by sonication (8 min, 30 sec on/off) and centrifugation (10,000 g, 10 min, 4°C), snap frozen and stored at −80°C. Western blot analysis was performed according to standard procedures using the following antibodies: anti-p21 (1:250, SXM30, BD Pharmingen), anti-Tubulin (1:5000, DM1A, Sigma-Aldrich). HRP-conjugated secondary antibodies (GE Healthcare) were used at a dilution of 1:5000 – 1: 7000).

### Cytokine array

Cytokine levels from tissue lysates were analyzed using the Mouse XL Cytokine Array kit (R&D Systems), following the manufacturer’s instructions. Cytokine arrays were developed using Invitrogen iBrightCL1000 imaging system (Thermo Fisher). Pixel intensity was determined using the ImageJ Software at different exposure times. The average background intensity of each membrane was subtracted prior to further analysis. The pixel intensity of each technical and biological replicate was then averaged for each condition. The final intensity values for each cytokine was arbitrarily set to 1 for the 2-3 months old mice and the values of the 6-8 months old mice were normalized accordingly. The final value is the mean of the normalized values at different exposure times.

### RT-qPCR and analysis

Frozen tissue was homogenized in TRI Reagent (MRC), purified using Direct-zol RNA Miniprep kit (Zymo Research) and reverse-transcribed using qScript cDNA Supermix (VWR International SAS). Values were normalized towards Actinb and Gapdh quantification. Real time qPCR was performed using gene-specific primers (Supplementary Table 1) and a LightCycler 480 (Roche).

### Statistical analysis

Statistical analysis was performed using the Prism 8 software (GraphPad Software Inc, San Diego, CA). Results are presented as mean ± S.E.M. Statistical significance was determined by the two-tailed unpaired Student’s *t*-test; *p ≤ 0.05, **p ≤ 0.01 and ***p ≤ 0.001.

## ACKNOWLEDGEMENTS

We thank Muriel Rhinn for help with procedures, and Matej Durik for comments on the manuscript. We thank the mouse facility at the Mouse Clinical Institute (ICS), the histology facilities at the CRG and the IGBMC/ICS, and the phenotyping platform at the ICS for excellent technical support. Work in the Keyes lab was funded in part by grants from: the Spanish Ministry for Economy and Competitiveness (SAF2013-49082-P), La Fondation Recherche Medicale (FRM) (AJE20160635985), Fondation ARC (PJA20181208104), IDEX Attractivité - University of Strasbourg (IDEX2017), and La Fondation Schlumberger pour l’Education et la Recherche (FSER) (FSER 19). Work was also supported by the grant ANR-10-LABX-0030-INRT, a French State fund managed by the Agence Nationale de la Recherche under the frame program Investissements d’Avenir ANR-10-IDEX-0002-02.”

---

**Supp. figure 1:**
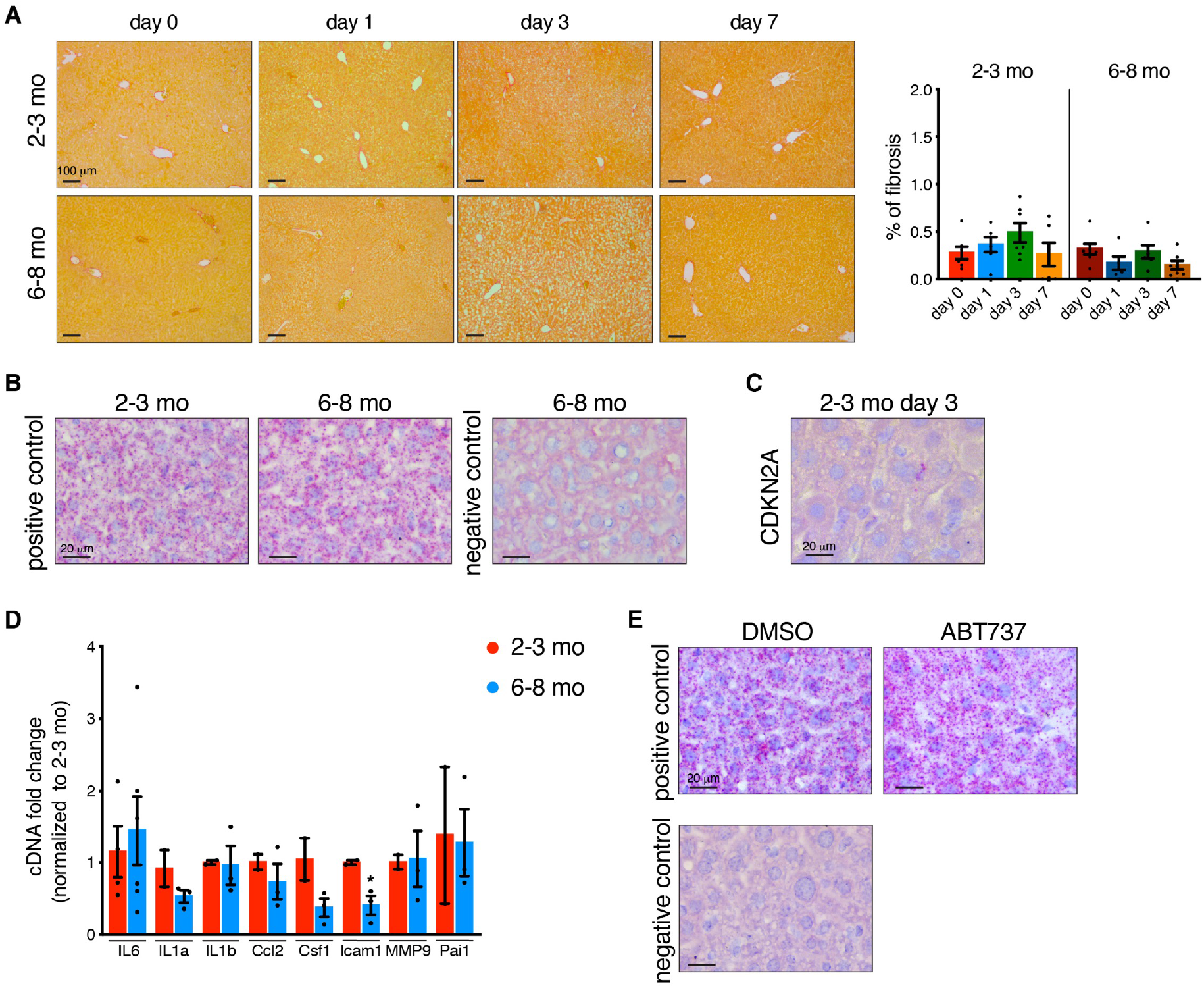
(A) Sirius red staining and quantification of young and adult liver sections before (day 0) and at different time points after PH. All images are representative of at least 5 biological replicates. Scale bars 100 μm. (B) RNA in situ hybridization staining for the positive control PPIB and bacterial DapB as negative control. All images are representative of at least 4 biological replicates. Scale bars 20 μm. (C) Representative image of RNA in situ hybridization staining for CDKN2A on 2-3 month old WT livers 3 days after PH (n=3). Scale bars 20 μm. (D) qPCR analysis for some SASP genes of 2-3 and 6-8 months old WT livers before PH (day 0) (n=2-6). (E) RNA in situ hybridization staining for the positive control PPIB and negative control bacterial DapB in DMSO and/or ABT-737-treated samples. All images are representative of at least 4 biological replicates. Scale bars 20 μm. Error bars, mean ± SEM, unpaired two-tailed Student’s *t*-test (*p ≤ 0.05, **p ≤ 0.01, ***p ≤ 0.001).

**Supp. figure 2:**
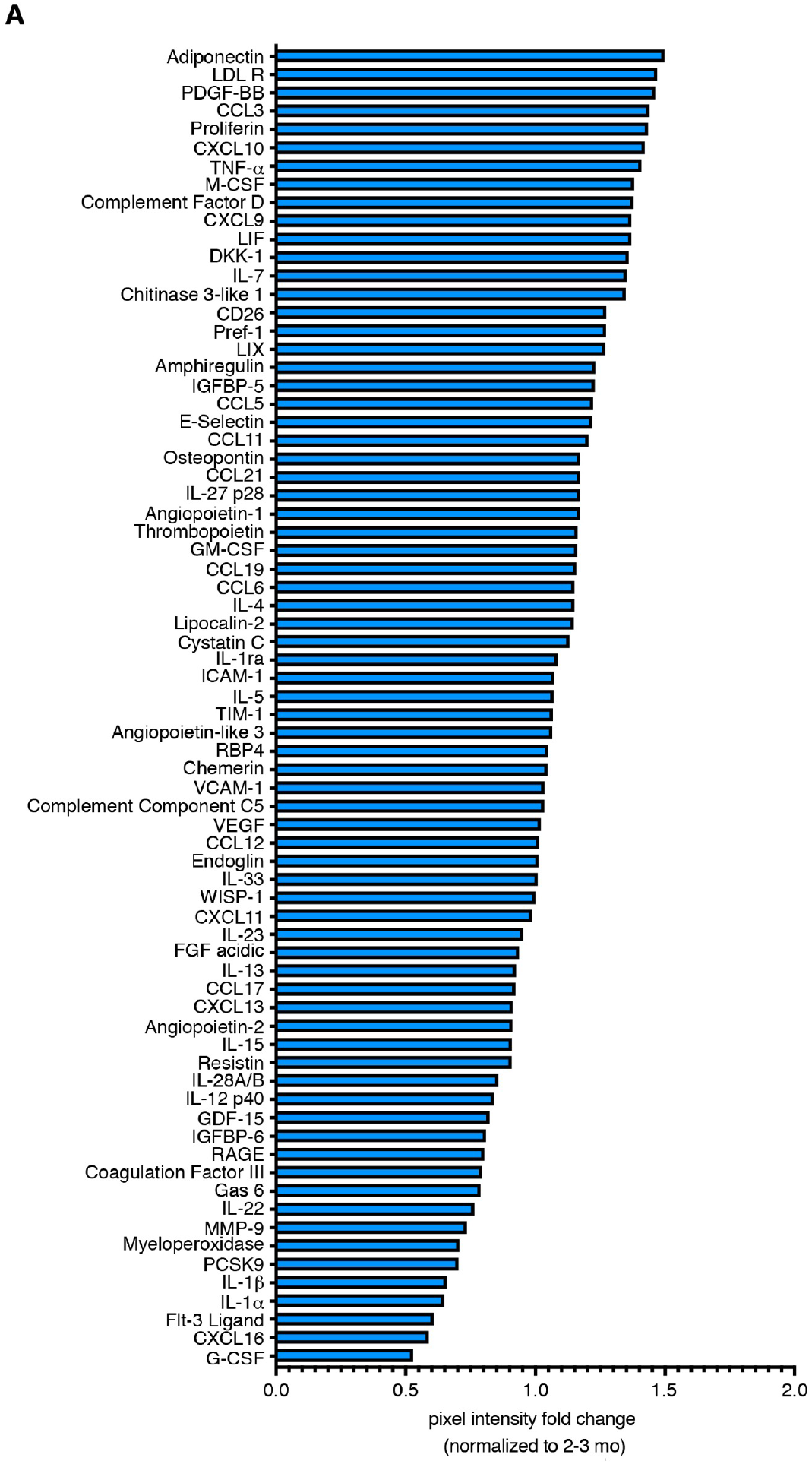
(A) Relative expression ratio (<1.5 fold) of 6-8 months old liver tissue 3 days after PH normalized to 2-3 months old livers (n=2).

**Supplementary Table 1.**
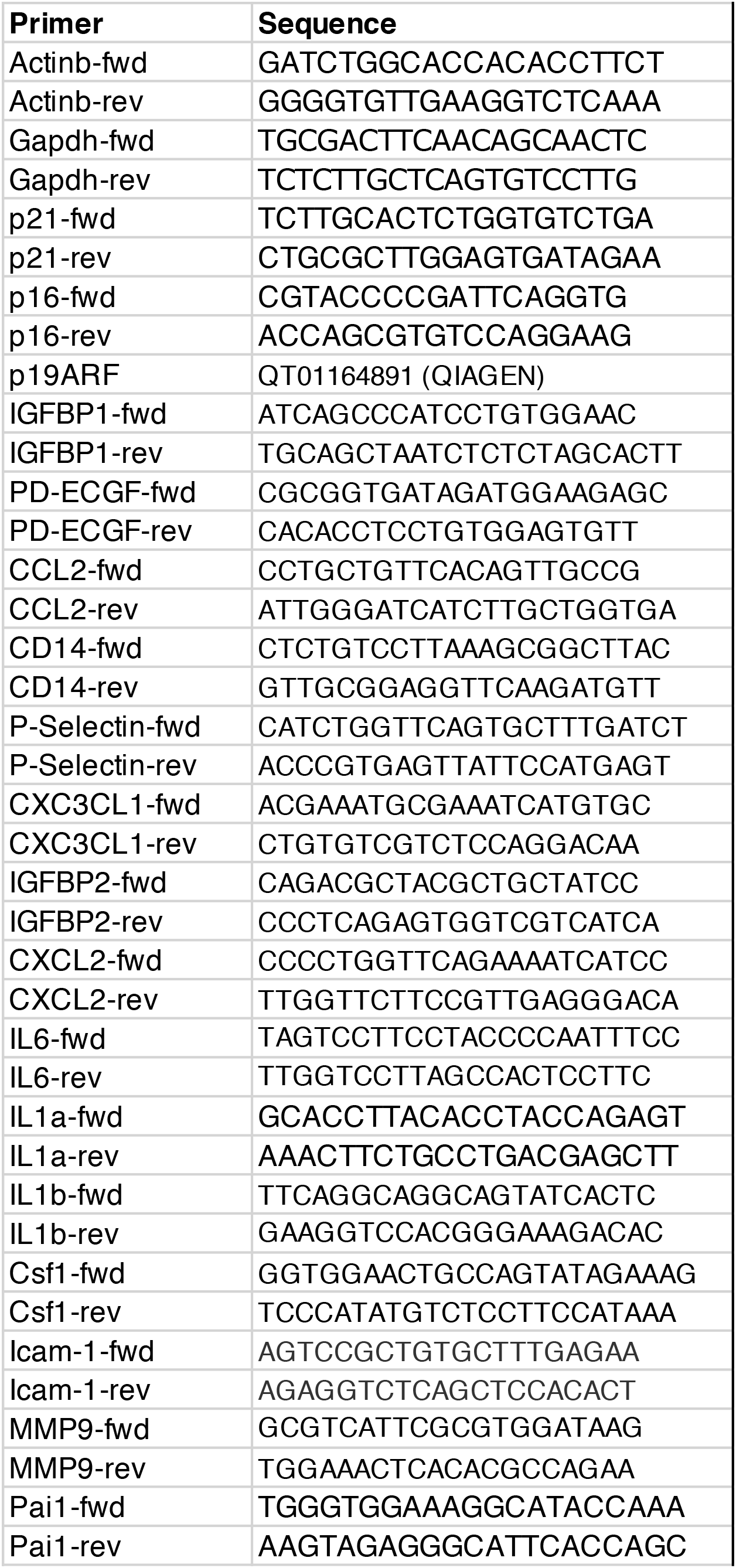
List of qPCR primers used

